# A shared wireless acquisition core for cortical and peripheral recordings in freely moving mice

**DOI:** 10.64898/2026.11.07.764907

**Authors:** Wanmei Zhang, Youlang Du, Jiayi Li, Guangyu Liang, Cheng Yi, Mingxiu Cheng, Xinhong Xv, Hao Wang, Hongyu Dong, Sen Song, Yuanlong Zhang, Zhiyuan Ma

## Abstract

Experiments in freely moving mice often require simultaneous recording of cortical, muscle and movement signals, but adding channels and peripheral sensors increases headstage burden and acquisition complexity. We developed an eight-channel wireless headstage that records six cortical channels and two neck-electrode channels, together with synchronized acceleration. Reassigning electrodes enables cardiac recording with the same electronics. The 13 × 11 × 3 mm board forms part of an approximately 1.2-g head-mounted assembly, with the battery carried separately on the back. With inputs internally shorted, root-mean-square noise was 0.176 μV over 0.5–100 Hz. In four mice with chemically induced seizures, recordings resolved distributed cortical activity and animal-level spectral changes. Subject-dependent evaluation of the seizure detector yielded an overlap-based event F1 of 0.893 ± 0.022. During pseudo-online replay, inference-software processing took 46.394 ± 0.363 ms per update. In a separate cohort, recordings revealed distinct coherence patterns associated with movement and social behavior, highlighting potential as biomarkers of behavior-related neural activity. Together, these results demonstrate concurrent cortical, muscle, and movement recording through a common acquisition chain, with electrode reassignment extending the same hardware to cardiac recording.

## Introduction

Interpreting cortical activity during natural behavior requires concurrent measurements of physiological and behavioral state^1,2,3^. Spontaneous and uninstructed movements can account for substantial variation in cortical activity, highlighting the importance of interpreting neural recordings alongside behavioral measurements. In freely moving mice, recording systems must limit head-mounted mass and characterize acquisition timing, transmission loss and data processing to support interpretation of the recordings^4,5^. Expanding sensor coverage or on-device computation can increase power requirements and headstage burden, creating a trade-off between recording capabilities and system demands.

Wireless multichannel recording and open acquisition software support neural measurements in freely moving animals^6,7,8^. Synchronized electrophysiological and behavioral monitoring^9^ and camera-coupled recording in freely moving mice^10^ enable joint analyses of neural activity and behavior. Recent studies have further advanced the quantitative characterization of social behavior through high-resolution tracking of postural dynamics and physical contact in freely interacting rodents. Electro-optic systems, skull-surface electrodes, flexible multiregional interfaces and multimodal platforms have expanded measurement capabilities and experimental control^11,12,13,14,15,16,17,18^, while open-source wireless rodent systems offer accessible implementations^19,20^. For experiments requiring distributed cortical signals together with muscle and motion measurements, a design challenge is to integrate these measurements into a compact acquisition chain that accommodates peripheral electrode reassignment and independent host-side display, storage and inference. Evaluating this design requires characterizing physiological coverage, headstage burden, and transmission performance across the intended recording configurations.

Here, we present a recording system that integrates an eight-channel headstage, a bilateral cortical and neck-electrode layout, synchronized acceleration, and independently buffered host-side processing. We characterize noise with shorted inputs and packet loss during battery-powered wireless operation, then examine cortical signals during pentylenetetrazol-induced seizures and natural behavior. Periodic visual stimulation tests frequency-specific signal content, and a separate electrode configuration evaluates cardiac recording with the same electronics. Subject-dependent seizure classification and pseudo-online replay assess the feasibility of host-side inference. Together, these experiments demonstrate flexible signal acquisition and compatibility with host-side analysis.

## Results

### The headstage integrates cortical, muscle, and motion recording

The headstage combines distributed cortical recording with neck-muscle and motion measurements on a single acquisition board. An eight-channel ADS1299 front end, a three-axis accelerometer, packetization, and a Bluetooth Low Energy (BLE) link were integrated on a 13 × 11 × 3 mm printed circuit board (Fig. 1a,b and Supplementary Fig. S1). The head-mounted assembly weighed approximately 1.2 g (excluding the battery) and included connectors, a flexible printed circuit (FPC), a machined housing, and a fixation base. An approximately 1.4-g battery was carried on the back, bringing the combined mass to approximately 2.6 g. Six channels recorded bilateral cortical signals, and the remaining two recorded signals from separate neck electrodes. Digital subtraction of the neck-electrode signals yielded one bipolar electromyography (EMG) signal. Three-axis acceleration was recorded within the same acquisition workflow. Thus, a single wireless system provided six cortical recordings, a derived bipolar EMG signal and three-axis acceleration measurements.

**Fig. 1.**
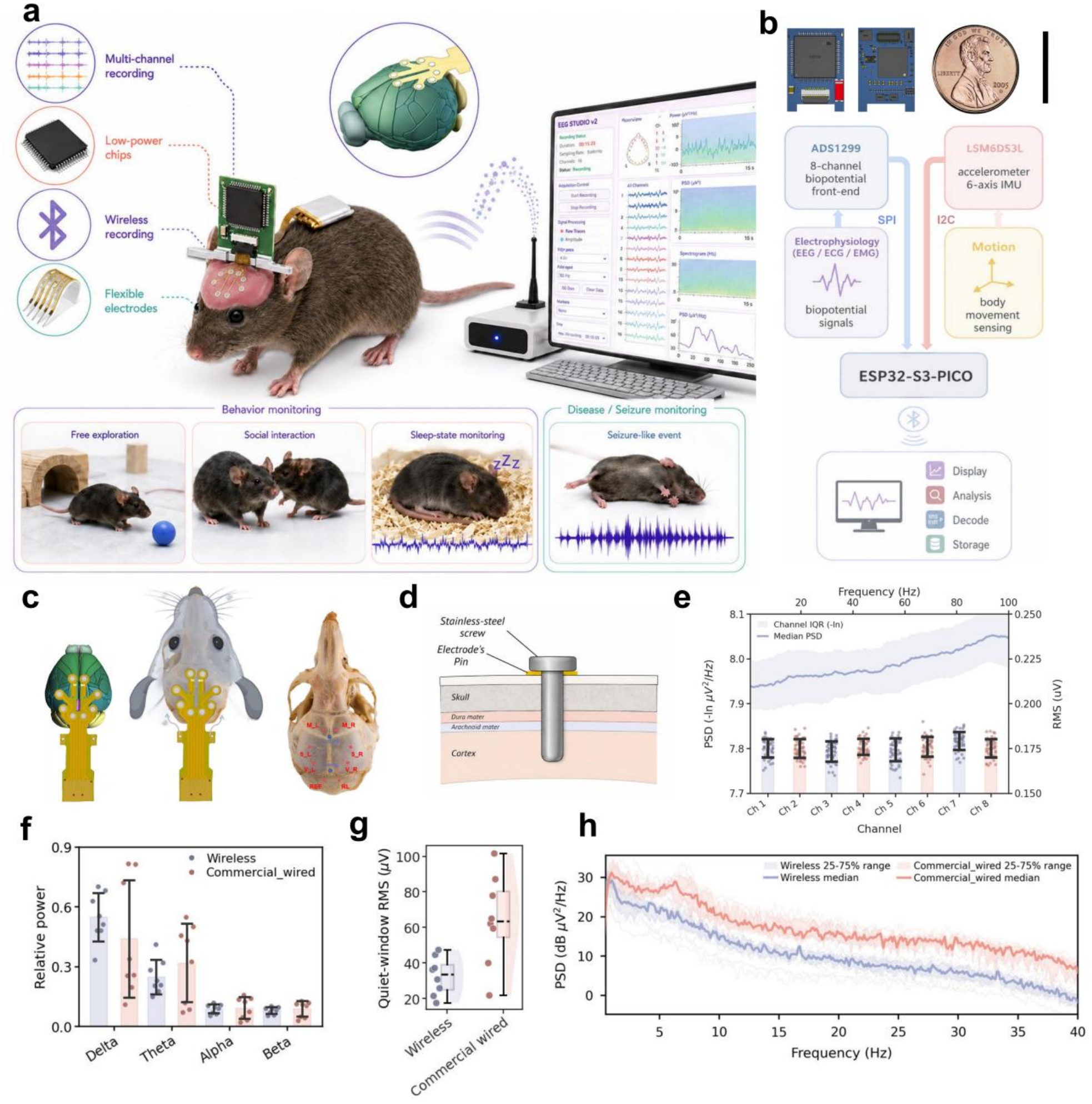
Lightweight wireless acquisition platform and signal characterization. a, Wireless headstage, flexible printed circuit (FPC), and graphical user interface. b, Architecture of the 13 × 11 × 3 mm printed circuit board and components of the approximately 1.2-g head-mounted assembly, with the approximately 1.4-g battery carried separately on the back. c, Bilateral cortical electrode sites over motor, somatosensory, and visual cortices, with reference and ground below lambda. d, Geometry of the 316L stainless-steel screw electrode. e, Input-short noise across eight channels during battery-powered Bluetooth Low Energy (BLE) transmission. The left axis shows the negative natural logarithm of PSD relative to 1 μV² Hz⁻¹, with shading indicating the across-channel interquartile range. The right axis shows 5-s window root-mean-square (RMS) summaries; the eight-channel mean full-recording RMS was 0.176 μV over 0.5–100 Hz. f, Relative cortical band power. g, Quiet-window RMS. h, Power spectral density before notch filtering. Panels f–h summarize eight recording sessions from seven mice per system, with separate cohorts for wireless and commercial recordings. Error bars and shading represent the quantities specified in the Methods.

A polyimide flexible printed circuit connected six 316L stainless-steel screw electrodes positioned over bilateral motor, somatosensory, and visual cortices (Fig. 1c,d). Each electrode was 0.5 mm in diameter with a 1.5-mm exposed length. The tips passed through the dura into the cortex, extending no more than 1 mm below the cortical surface. Reference and ground electrodes were placed below lambda over the cerebellum (Supplementary Fig. S2). This bilateral layout enabled site-resolved comparisons within the same recording, demonstrating the importance of electrode placement beyond channel count alone.

During battery-powered BLE transmission with all eight inputs internally shorted and gain set to 24, mean full-recording input-referred root-mean-square (RMS) noise was 0.176 μV over 0.5–100 Hz (Fig. 1e and Supplementary Fig. S3). Across approximately 30 experiments at distances up to 5 m, mean packet loss was approximately 2% at a sampling rate of 1,000 Hz and 0.2% at 250 Hz. These results characterize system performance under the tested conditions. Recording duration and power consumption were not systematically benchmarked.

We also compared the amplitudes and spectra of physiological signals recorded wirelessly and with a PowerLab 2/26 recording set (Fig. 1f–h). Relative band-power distributions were broadly similar. Median RMS amplitude during quiet windows was 33.74 μV for wireless recordings and 63.66 μV for PowerLab recordings.

### A unified host platform integrates monitoring, recording and seizure detection

We developed a host-side platform that integrates visualization, continuous recording, and seizure detection, with the headstage dedicated to signal acquisition and wireless transmission (Fig. 2a,b). After packet checking and decoding, incoming data were routed to three independently buffered branches for display, asynchronous HDF5 storage and inference. The interface displayed 10-s cortical traces, spectrograms, differential neck-EMG RMS, and three-axis acceleration. The detection branch used a rolling 30-s history of six cortical channels to update seizure probability once per second.

**Fig. 2.**
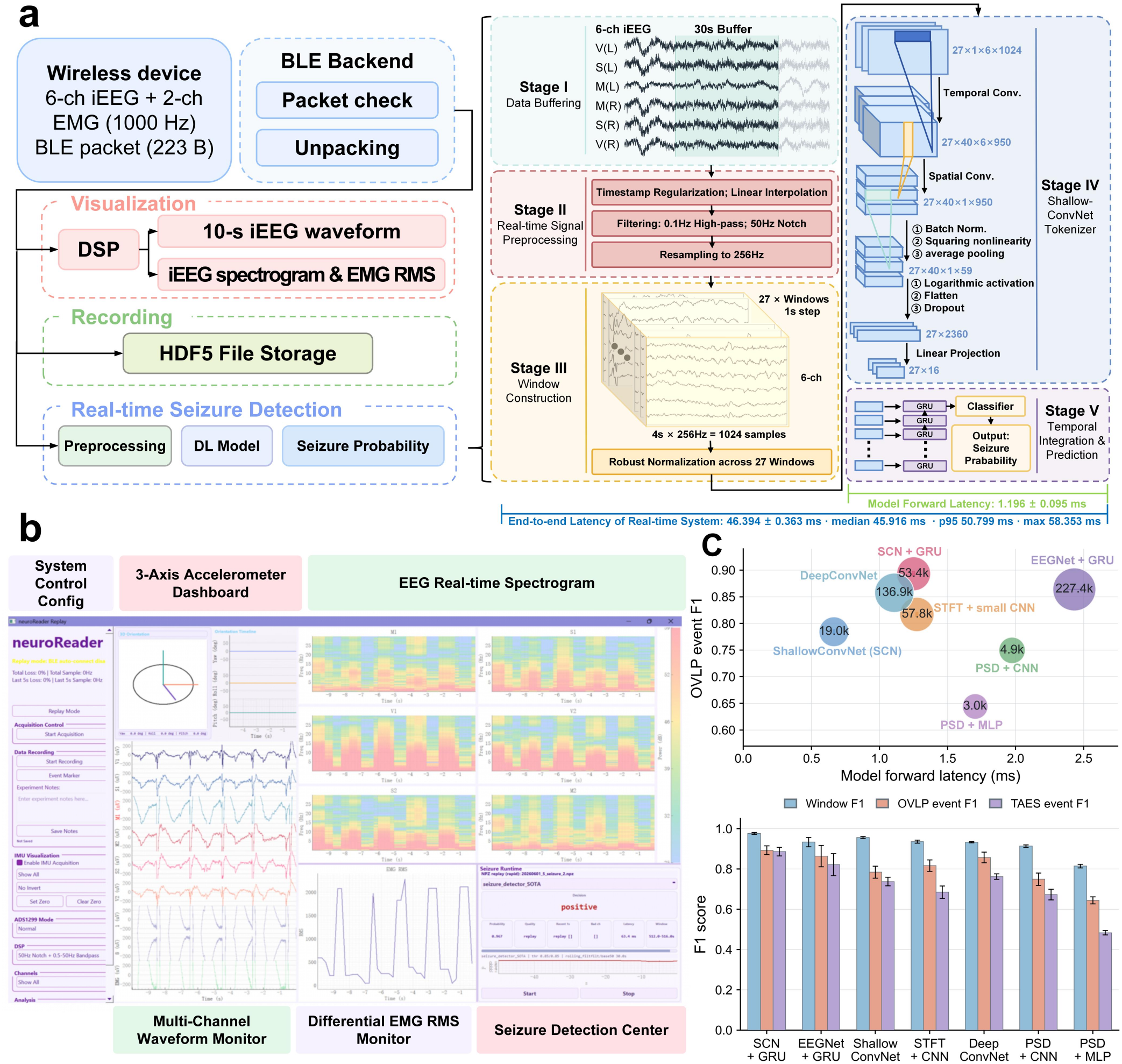
Modular host workflow for streaming acquisition and online seizure detection. a, Workflow showing BLE reception, packet processing, visualization, asynchronous HDF5 storage, and seizure detection. The selected ShallowConvNet–GRU contains 53,434 trainable parameters. Model forward-pass latency and inference-software processing time are shown, with timing boundaries defined in the Methods. b, Graphical user interface displaying system configuration, three-axis acceleration, multichannel cortical signals, spectrograms, differential neck-EMG RMS, storage status, and seizure estimates. c, Performance and efficiency of seven candidate models. The upper panel plots overlap-based event F1 against model forward-pass latency measured with a batch size of one and GPU-resident inputs; bubble size indicates parameter count. The lower panel shows window-level F1, overlap-based event F1, and time-aligned event F1. Bars indicate means, and error bars indicate SDs across 10 random seeds.

To identify a suitable detector, we compared seven candidate models using a subject-dependent split and 10 training seeds (Fig. 2c and Supplementary Fig. S4). The selected ShallowConvNet–GRU combined 27 overlapping 4-s windows within each 30-s context to incorporate recent signal history, with 53,434 trainable parameters. It achieved a window-level F1 of 0.977 ± 0.005 and an overlap-based (OVLP) event F1 of 0.893 ± 0.022.

Across three HDF5 pseudo-online replay runs on an NVIDIA RTX 4090, the pipeline completed processing well within the 1-s update interval. Model forward-pass latency was 1.196 ± 0.095 ms, and software processing from an available input chunk to the prediction output took 46.394 ± 0.363 ms per update. Together, these results demonstrate the integration of multichannel monitoring, continuous recording, and computationally efficient seizure-probability updates within a unified host-side workflow.

### Multisite recordings capture seizure-associated patterns

We examined state-dependent cortical activity across six recording sites in four eight-week-old male wild-type mice before and after pentylenetetrazol administration (PTZ; 40 mg kg^−1^; Fig. 3a). Six-channel waveforms showed recurrent rhythmic epileptiform discharges, and Morlet time–frequency maps revealed transient power elevations (Fig. 3b). The headstage thus captured temporally structured seizure-associated signals across the implanted sites.

**Fig. 3.**
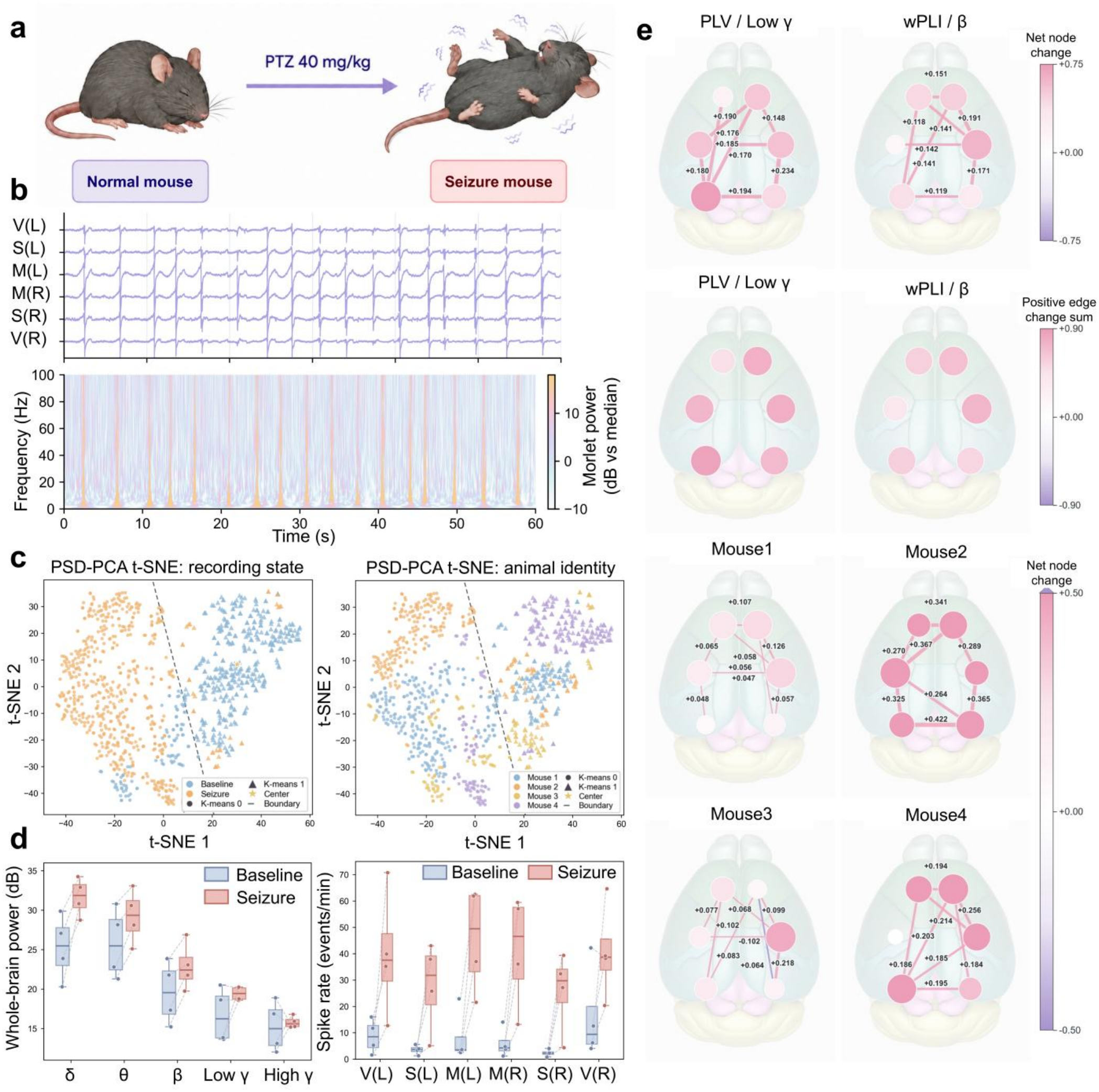
Cortical recording and multiscale analysis during PTZ-induced seizure activity. a, Recording paradigm before and after PTZ administration (40 mg kg⁻¹; n = 4 mice). b, Representative 60-s six-channel recording and corresponding Morlet time–frequency map. c, PCA–t-SNE embedding of 1,035 overlapping 10-s windows from 13 accepted segments, colored by animal identity or physiological state. d, Animal-level band power summarized across cortical channels and sharp-transient rates at individual recording sites. Each point represents one mouse, with lines connecting paired observations across conditions. Boxes indicate the interquartile range (IQR), center lines the median, and whiskers extend to the most extreme observations within 1.5 × IQR of the quartiles. e, Seizure-minus-baseline changes in low-gamma phase-locking value (PLV) and beta-band weighted phase-lag index (wPLI), showing group summaries and individual beta-band wPLI maps. Displayed edges were selected by effect magnitude rather than statistical significance.

Spectral features showed separation between baseline and seizure windows in the principal component analysis–t-distributed stochastic neighbor embedding (PCA–t-SNE) visualization, with variation across animals (Fig. 3c). Delta-, theta- and beta-band power increased in all four mice, whereas gamma-band changes varied across animals (Fig. 3d). Sharp-transient rates increased at five cortical sites in all four animals and at the right visual site in three animals. Together, spectral and transient-event measures captured complementary features of the transition from baseline to seizure activity.

Multisite recording further revealed seizure-associated changes in phase coupling across cortical regions^21,22^. We quantified coupling across all 15 cortical pairs using phase-locking value (PLV)^23^ and weighted phase-lag index (wPLI)^24^. Group-level maps showed increased low-gamma PLV and beta-band wPLI during seizures, while individual maps revealed animal-specific spatial patterns (Fig. 3e and Supplementary Fig. S5). These connectivity maps complemented power and transient-rate measurements by revealing how seizure-associated changes were organized across cortical regions. Together, these analyses provided a multiscale characterization of seizure activity, from individual recording sites to distributed networks.

### Wireless multisite recording enables cortical network analysis in freely moving mice

We next evaluated the platform in two freely moving behavioral settings by combining multisite cortical recordings with video-based behavioral tracking. During exploration of a 40 × 40 cm open field, overhead video provided trajectories and speed-based movement labels for comparison with cortical activity^25^ (Fig. 4a–c). In four mice, coherence differences between moving and still periods varied across cortical connections and frequency bands, with a subset remaining significant after Benjamini–Hochberg correction (Fig. 4d). These analyses characterized the spatial and spectral organization of movement-associated cortical coupling, showing how wireless multisite recordings can extend behavioral analysis beyond trajectories and locomotor measures. Locomotor measurements with and without the head-mounted device are compared in Supplementary Fig. S7.

**Fig. 4.**
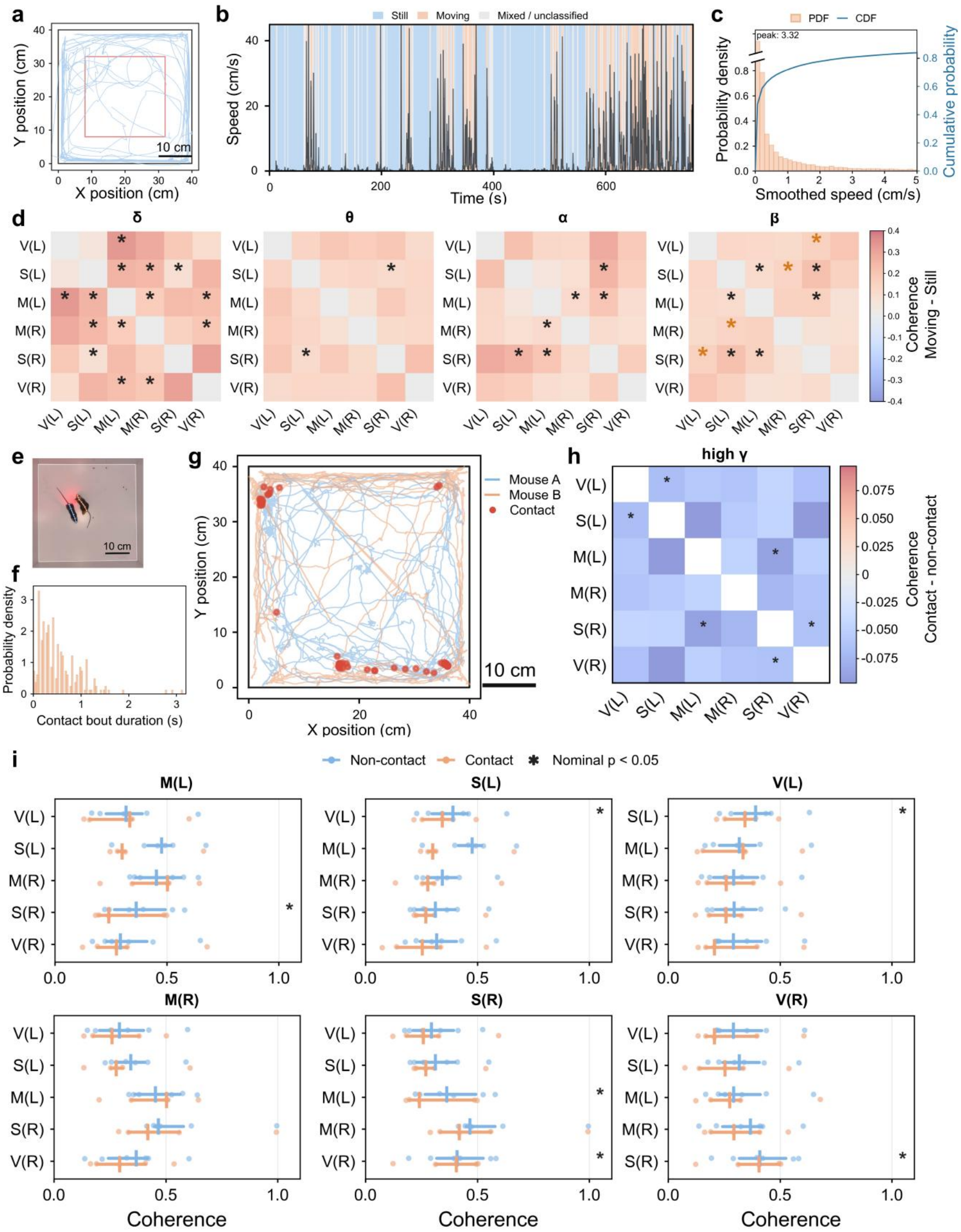
Behavioral tracking and cortical coupling during free exploration and social contact. a, Representative trajectory in a 40 × 40 cm open field. b, Smoothed speed and movement-state labels (still, moving, and mixed). c, Probability density and cumulative distribution of speed. d, Animal-level moving-minus-still coherence differences in delta, theta, alpha, and beta bands (n = 4 mice). Black asterisks indicate uncorrected P < 0.05; orange asterisks indicate Benjamini–Hochberg-adjusted P < 0.05. e, SLEAP pose estimates for two animals. f, Distribution of detected contact-bout durations. g, Representative trajectories and contact-associated locations. h, Contact-minus-non-contact differences in the high-gamma spectral coherence index, averaged across animals. i, Seed profiles of the same index during contact and non-contact periods (n = 5 mice).

We extended this approach to social-contact experiments, using SLEAP tracking of two animals to identify contact bouts and map their locations within the arena (Fig. 4e–g). We aligned these annotations with cortical recordings and compared the short-window, bandwise spectral coherence index between contact and non-contact periods for each cortical connection. In five mice, connectivity maps and seed profiles revealed variation across cortical connections and animals (Fig. 4h,i). Some connections showed nominally significant contact-associated differences; none remained significant after Benjamini–Hochberg correction at the animal level in the primary analysis (Supplementary Fig. S6). Together, the exploration and social-contact experiments demonstrate an integrated workflow linking wireless recording and behavioral tracking to region- and frequency-resolved analyses of cortical coupling, highlighting the platform’s ability to investigate neural activity in freely moving mice.

### Wireless cortical recording resolves periodic activity during visual stimulation

We next evaluated the ability of the wireless platform to resolve frequency-specific cortical activity during periodic visual stimulation^26^ (Fig. 5a). Recordings from the right visual cortex contained a prominent spectral component near 5.25 Hz during nominal 5-Hz stimulation. Using this component as a reference, we divided a continuous recording excerpt into 80 non-overlapping windows, each spanning five reference cycles. Overlaid windows showed variation in individual waveforms, and their average retained a recurring oscillatory pattern (Fig. 5b). This provided a time-domain characterization of rhythmic cortical activity that complemented the spectral analysis.

**Fig. 5.**
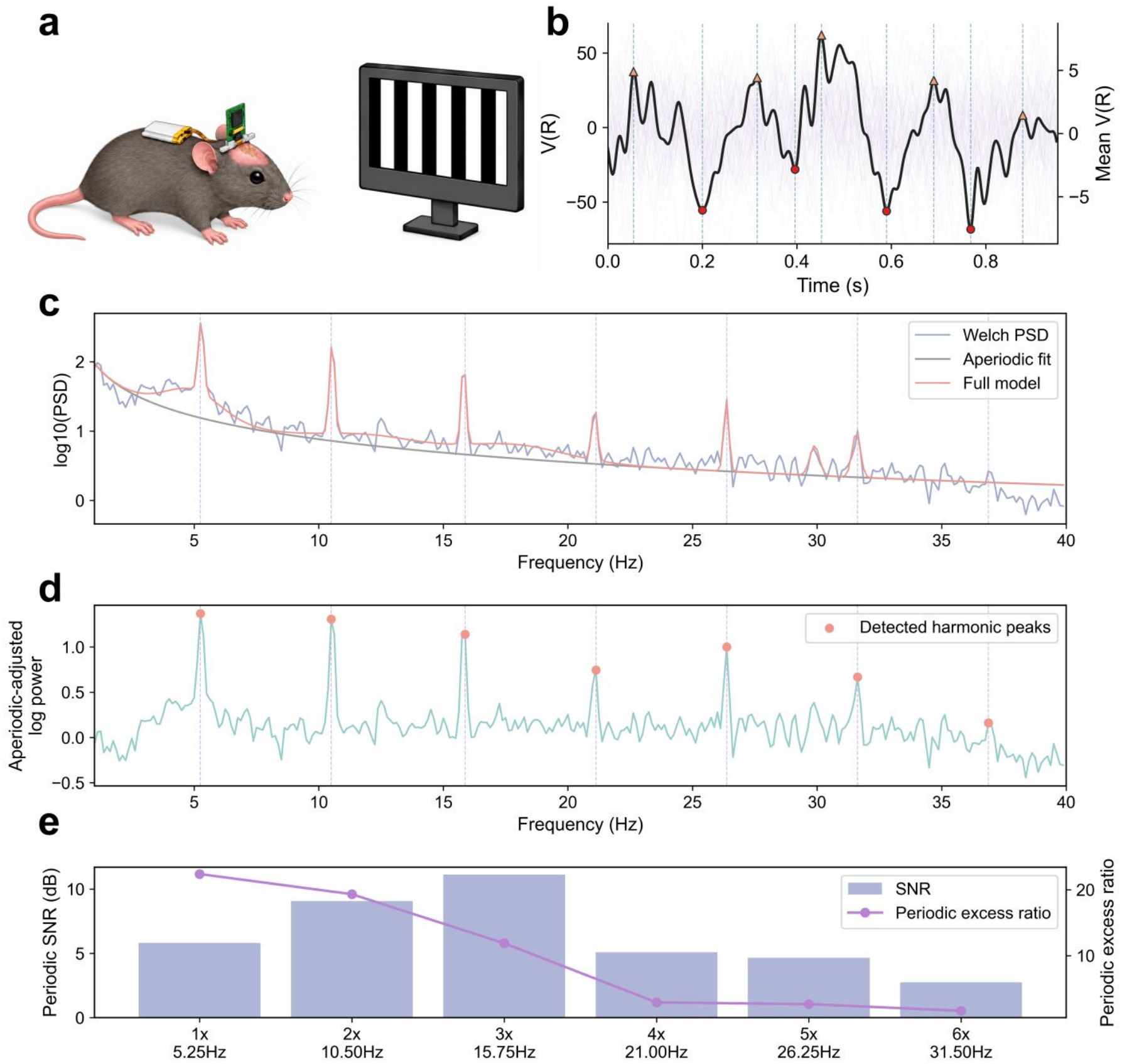
Frequency-specific cortical recording during visual stimulation. a, Visual-stimulation arrangement. b, Waveforms recorded from the right visual cortex, divided into 80 non-overlapping windows, each spanning five cycles at the 5.25-Hz reference frequency. The thick line indicates the mean. c, Welch power spectral density with fitted aperiodic and full spectral models. d, Aperiodic-adjusted log power obtained by subtracting the fitted aperiodic log spectrum. e, Local harmonic signal-to-noise ratio (left axis) and periodic excess ratio relative to the fitted aperiodic background (right axis). Panels b–e use the same recording: b shows a selected continuous excerpt, and c–e summarize the full preprocessed recording.

Welch power spectra and spectral parameterization identified components near the 5.25-Hz reference and its harmonics above the fitted aperiodic background^27^ (Fig. 5c). Removing this background clarified the periodic components across the spectrum (Fig. 5d). Periodic excess ratios quantified their elevation above the fitted background, and local signal-to-noise estimates measured their contrast with neighboring frequencies (Fig. 5e). Together, these waveform and spectral analyses demonstrate that the wireless platform can capture rhythmic cortical activity and resolve its harmonic structure, extending its application from spontaneous behavioral recordings to controlled sensory-stimulation experiments.

### Electrode reconfiguration extends the wireless platform to cardiac recording

Finally, we demonstrated the reconfigurability of the platform by adapting it for electrocardiography (ECG), while retaining the same acquisition board and wireless workflow. Electrodes placed subcutaneously near the right forelimb and left hindlimb provided a configuration for cardiac recording (Fig. 6a). Recordings showed recurring cardiac complexes with prominent negative-going QRS deflections. Alignment to the detected fiducial points revealed a consistent waveform across cardiac cycles (Fig. 6b,c). We measured beat-to-beat intervals between successive detected QRS complexes and used these intervals to calculate instantaneous heart rate (Fig. 6d,e). Together, these results demonstrate the use of the wireless platform for cortical, muscle and cardiac recordings through different electrode configurations while retaining the same acquisition electronics.

**Fig. 6.**
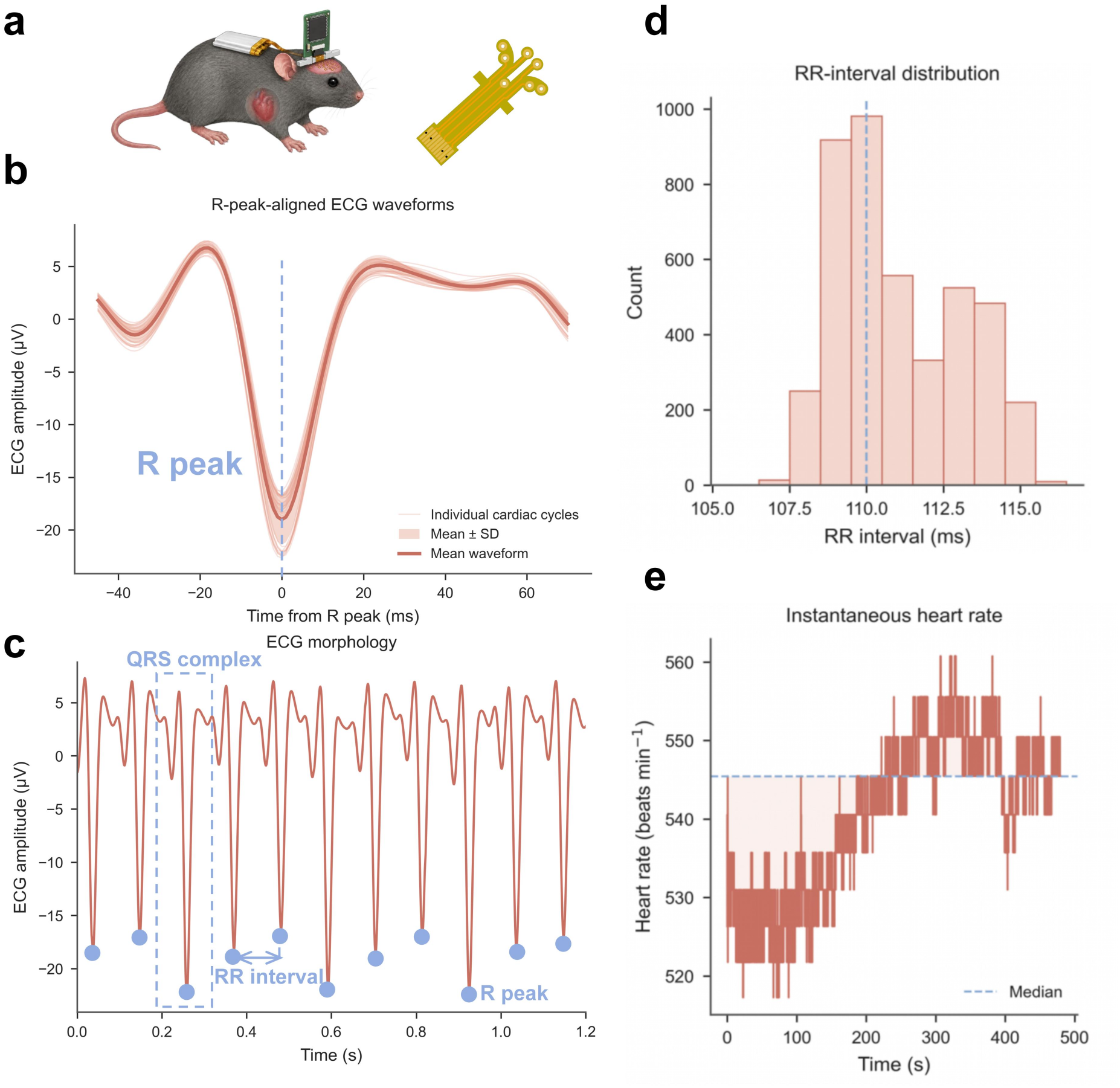
Peripheral electrode reassignment enables wireless cardiac recording. a, ECG electrode configuration using the same acquisition electronics. b, Cardiac waveforms aligned to detected negative-going QRS fiducial points. The thick line indicates the mean, and the shaded area indicates mean ± SD. c, Representative continuous ECG trace. d, Distribution of beat-to-beat intervals. e, Instantaneous heart rate calculated from intervals between consecutive detected beats.

## Discussion

This study presents a compact wireless acquisition platform that combines configurable electrode arrangements with host-side processing to support physiological recording and analysis. Hardware characterization in Fig. 1 establishes the foundation for this approach: the eight-channel acquisition board forms part of an approximately 1.2-g head-mounted assembly, excluding the battery, and achieved an input-referred noise of 0.176 μV RMS over 0.5–100 Hz with internally shorted inputs during wireless operation. Carrying the approximately 1.4-g battery on the back keeps energy storage off the head and distributes the combined mass of approximately 2.6 g between the head and back. Table 1 compares this arrangement with battery-integrated headstages in several comparator systems and places the device dimensions, channel coverage and recording performance in context. This architecture provides a common acquisition core whose physiological coverage and analysis functions can be adapted through electrode reconfiguration and host-side processing.

**Table 1.** Comparison of recording hardware for small-animal electrophysiology.

| **Metric** | **Cheng & Murari, 2020<sup>51</sup><br>OSERR** | **Kim et al., 2020<sup>52,53</sup><br>CBRAIN;<br>EEG+EMG** | **Lee et al., 2023<sup>54</sup><br>Wireless ECoG** | **Benomar et al., 2024<sup>55</sup><br>Wireless mouse EEG<br>(EMBC)** | **Lim et al., 2024<sup>56</sup><br>NAT-1 application** | **Shin et al., 2025<sup>57</sup><br>NeuroStim** | **Pinnacle<br>(commercial)<sup>58</sup><br>Wireless mouse<br>EEG/EMG** | **This work<br>Flexible wireless<br>iEEG/EMG** |
|---|---|---|---|---|---|---|---|---|
| Animal model | Mouse | Mouse | Rabbit | Mouse | Mouse | Rat | Mouse | **Mouse** |
| Module dimensions (mm) | 13 × 8 × 8 | 16 × 28 × 6 | 18 × 15 × 10 | Ø 21.5 × 9.8 | 18 × 22.2 × 10.3 | 16 × 28 × 6 | 17.1 × 16.2 × 11.0 | **13 × 11 × 3** |
| Reported device mass (g) | 1.8 (incl. battery) | 2.6 (incl. battery) | 2.1 (electronics);<br>9.5 (incl. battery + enclosure) | ≥3.79 (incl. battery) | 2.2 (incl. battery);<br>~3.4 (incl. mount) | 3.8 (incl. battery) | 3.8 (incl. battery) | **~1.2 (head)**<br>**~1.4 (back battery)**<br>**~2.6 (combined)** |
| Battery location | Head | Head | — | — | Head | Head | Head | **Back** |
| Battery capacity (mAh) | 30 | 40 | — | — | — | 150 | — | **100 / 150** |
| Neural recording channels | 2 differential | 16-ch hardware capability | 32 ECoG + 4 reference | 4 EEG | 4 EEG | 16 recording + 16 stimulation | 2 EEG + 1 EMG; or 3 EEG | **6 iEEG + 2 neck inputs (1 derived EMG)** |
| Sampling rate (S/s/ch) | 240 | 1,024 (in vivo);<br>256–16,000 (settable) | 6,000 (in vivo);<br>30,000 (max) | 280 | 250–2,000 | 512 | 256 / 512 / 1,024 | **250 / 500 / 1,000** |
| ADC resolution (bit) | 10 | 16 | 16 | 24 | 12 (11-bit signed output) | 16 | 12 | **24 (ADS1299)** |
| Input-referred noise (µV RMS) | 2.3 (shorted inputs) | — | — | — | — | — | — | **0.176 (shorted inputs;<br>0.5–100 Hz)** |
| Wireless / data path | On-board memory;<br>magnetic start | 2.4-GHz ESB | 2.4-GHz Wi-Fi / UDP | BLE | On-board memory; IR event stamps | BLE 4.0 | Bluetooth 5 | **BLE** |
| Wireless range (m) | N/A | 2 | — | — | N/A | 6 | ≥3.0 | **5** |

Notes. —, unavailable; N/A, not applicable. Mass includes the battery, enclosure or mount only where indicated. Battery location is listed where established by the cited documentation. In this work, approximately 1.2 g is carried on the head and the approximately 1.4-g battery is carried on the back, giving a combined mass of approximately 2.6 g. The head-mounted mass is therefore distinct from the total mass. Recording channels and stimulation channels are listed separately; two neck-electrode inputs yield one derived bipolar EMG channel in this work.
Abbreviations: ADC, analog-to-digital converter; BLE, Bluetooth Low Energy; ECoG, electrocorticography; EEG, electroencephalography; EMG, electromyography; ESB, Enhanced ShockBurst; iEEG, intracranial EEG; IR, infrared; RMS, root mean square; S/s/ch, samples per second per channel; UDP, User Datagram Protocol.

Offline analyses demonstrate that multisite recordings support complementary measurements of local cortical activity and coupling across regions. During PTZ-induced seizures, the six-channel layout captured site-specific spectral and sharp-transient changes alongside phase coupling across cortical pairs. These measures describe both local electrophysiological changes and the spatial organization of seizure-associated cortical coupling. Recordings during visual stimulation further resolved periodic components and harmonic structure above the fitted aperiodic background. Together, these findings establish a common recording framework for waveform, spectral and connectivity analyses.

Free-exploration and social-contact experiments further demonstrate the platform’s applicability to complex behavioral settings. Removing the recording cable reduces mechanical constraints during arena exploration and social interaction^28^. Combining wireless cortical recordings with video tracking linked neural measurements to trajectories, movement states and contact bouts^29^. The multisite layout enabled behavioral comparisons of coupling across connections and frequency bands, complementing individual-channel measurements. These demonstrations establish an experimental workflow integrating behavioral tracking in freely moving mice with cortical network analysis, providing a framework for examining how distributed cortical activity varies during exploration and social interaction.

The ECG experiment further demonstrates the platform’s adaptability across physiological recording applications. By reconfiguring electrode placement, the same acquisition electronics and wireless workflow supported cardiac waveform recording and beat-to-beat timing analysis, extending the platform’s physiological scope without redesigning the core electronics. This shared multichannel architecture provides a basis for future configurations that allocate channels to neural and peripheral electrical signals, supporting the development of simultaneous multimodal electrophysiological recording in small animals.

Wireless rodent systems expand experimental access through open designs, multimodal integration, and autonomous operation. Open-source electrophysiology platforms support untethered recording and adaptation of acquisition hardware and software^19,20^. WILD integrates neural and behavioral sensors with onboard event processing and closed-loop manipulation^15^, and battery-free implants combine physiological recording with autonomous neuromodulation^30^. Our platform pairs a compact, reusable acquisition core with configurable electrodes and adaptable host-side processing. Electrode reconfiguration and host-side analysis allow recording targets and analytical tasks to be adapted without redesigning the acquisition board. This flexibility supports reuse of hardware and acquisition procedures across studies combining distributed cortical measurements with behavioral or peripheral physiological information.

The GUI and seizure-detection experiments demonstrate a host-side workflow that combines continuous recording, interactive monitoring, and regularly updated predictions. Placing visualization, storage, and inference on the host computer keeps the headstage dedicated to acquisition and transmission and allows analysis software to evolve independently of the recording electronics. Within-recording model evaluation and HDF5 pseudo-online replay demonstrated the integration of seizure detection into this workflow, with software processing completed within the 1-s update interval during replay. Integrating stimulation control could extend this architecture to wireless closed-loop experiments in which detected neural events guide intervention^31,32^. Building on studies across experimental epilepsy models^33^ and long-duration mouse recordings^34^, prospective testing in independent animals would quantify detection accuracy, false-alarm rates, and response timing required for that application^35,36^.

Further development should extend the operating range, improve wireless continuity at higher sampling rates, and strengthen validation over longer recordings. Mean packet loss increased from approximately 0.2% at 250 Hz to 2% at 1,000 Hz, identifying wireless continuity at higher data rates as an engineering priority. Longer recordings would characterize battery endurance and electrode stability^37^. Comparisons with commercial systems currently involve separate cohorts and channel configurations; matched recordings would provide a more direct assessment of recording performance.

Larger biological studies would assess the consistency of movement- and social-contact-associated coherence patterns across animals. Some movement-associated differences remained significant after multiple-comparison correction, whereas social-contact differences did not survive correction at the animal level. Recording stimulus triggers would enable stimulus-aligned analyses of visual responses, and wearable mouse ECG systems^38^ provide a reference for future cardiac-specific validation. These studies would define the platform’s long-term performance, temporal precision, and applicability across animals and recording conditions.

Overall, the platform integrates multisite recording, reconfigurable physiological interfaces, and adaptable host-side analysis in a common experimental workflow. Hardware characterization and demonstrations in seizure monitoring, freely moving behavior, sensory stimulation, and cardiac recording establish its use across these experimental settings. Electrode placement and software analysis adapt the same wireless acquisition hardware to different neural and peripheral physiological recording applications.

## Methods

Supplementary Methods S1–S5 describe branch-specific preprocessing, detector implementation and scoring, and connectivity, behavioral, and software settings. Supplementary Table S1 lists analysis-specific filter settings.

### Animals and electrode implantation

PTZ experiments used four eight-week-old male wild-type mice. Wireless and commercial recording comparisons used separate cohorts of seven mice each, with eight recording sessions per system. Movement-state and social-contact analyses included four and five mice, respectively, and one mouse participated in both analyses.

A polyimide flexible printed circuit (FPC) connected six 316L stainless-steel screw electrodes to the headstage. Each electrode had a diameter of 0.5 mm and an exposed length of 1.5 mm. Bilateral recording sites were positioned relative to bregma over motor cortex (AP +1.5 mm, ML ±1.0 mm), somatosensory cortex (AP −1.0 mm, ML ±3.2 mm), and visual cortex (AP −4.5 mm, ML ±2.5 mm). Reference and ground contacts were placed below lambda over the cerebellar region. Signals from two neck electrodes were acquired through separate channels and subtracted in software to derive one bipolar EMG signal.

Electrode continuity was checked before implantation. Under isoflurane anesthesia, mice were secured in a stereotaxic frame, the skull was exposed, and drilling sites were identified. Screw electrodes were inserted through the skull and dura, with their tips extending into the cortex no more than 1 mm below the cortical surface. This insertion depth was distinct from the 1.5-mm exposed screw length. Dental cement and the machined housing with its implantation fixation base secured the assembly.

### Ethics statement

All animal procedures were approved by the Institutional Animal Care and Use Committee (IACUC) of Tsinghua University and were performed in accordance with relevant guidelines and regulations.

### Wireless hardware and data acquisition

The headstage integrated an eight-channel ADS1299 analog front end, a three-axis accelerometer, and an ESP32-S3-PICO controller on a 13 × 11 × 3 mm PCB. The approximately 1.2-g head-mounted assembly included the PCB, connectors, FPC, machined housing, and implantation fixation base, excluding the battery. The approximately 1.4-g battery was mounted on the back, bringing the combined headstage-and-battery mass to approximately 2.6 g. Experiments used nominal battery capacities of 100 or 150 mAh. Six channels acquired cortical signals, and two acquired the individual neck-electrode signals, which were subtracted in software to derive one differential EMG signal. Biopotentials were acquired at a gain of 24, sampled at 1,000 or 250 Hz, and transmitted over BLE with device timestamps.

The desktop application routed received samples to visualization, asynchronous HDF5 storage, and seizure inference. Displays included cortical traces, spectrograms, differential EMG RMS, and three-axis acceleration. The deployed ShallowConvNet–GRU used the six cortical channels as input. The other six benchmark models additionally used the differential neck-EMG signal according to their respective model configurations. Transmission was tested across approximately 30 experiments at distances up to 5 m. Packet loss was calculated as the number of missing packets divided by the number of expected packets. Mean packet loss was approximately 2% at 1,000 Hz and 0.2% at 250 Hz. These values represent averages across the tested experiments rather than maximum packet-loss rates.

### Signal preprocessing and recording validation

Preprocessing followed analysis-specific workflows. Where timestamp reconstruction was used, recordings were screened for invalid timestamps and non-finite samples, ordered by timestamp, and deduplicated. The native sampling rate was estimated from the median positive timestamp increment. Signals were then interpolated onto a uniform time grid and demeaned channel by channel. Input-short, visual-stimulation, and ECG analyses used uniformly sampled sequences without timestamp reconstruction. The 0.5–40 Hz passband used for commercial recordings was not applied to separate analyses of higher-frequency wireless signals.

Electronic noise was measured with all eight ADS1299 inputs internally shorted at gain 24 during battery-powered BLE transmission. The 1-kHz record was treated as a uniform sample sequence, and Welch power spectral density (PSD) estimates were summarized across channels^39^. For RMS calculations, signals were filtered over 0.5–100 Hz using a fourth-order Butterworth filter applied bidirectionally. Full-recording channel RMS and the arithmetic mean across eight channels were calculated as:

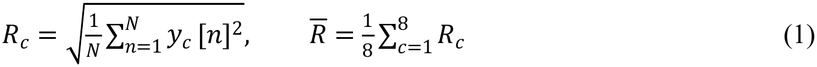

Here *y_c_*[*n*] is the filtered input-referred voltage at sample n in channel c, and N is the number of samples. Full-recording RMS included all filtered samples. Separately, RMS values from non-overlapping 5-s windows were screened using within-channel 1.5 × IQR limits and summarized as mean ± SD. Outlier exclusion applied only to these window-level summaries.

Wireless and PowerLab 2/26 recordings were acquired in separate, unpaired sessions. Each wireless session included six cortical channels, and each commercial session included one EEG channel. Recordings were divided into non-overlapping 10-s windows and screened for excessive amplitudes or first-order temporal differences, flat signals and narrowband contamination. Accepted windows were used to calculate PSD before notch filtering and relative band power and quiet-window RMS after notch filtering. Relative band power was defined as the PSD integral over each frequency band divided by the integral over 0.5–40 Hz.

Within each channel, the 20% of accepted windows with the lowest RMS were designated as quiet windows based on RMS alone, without using behavioral labels. Wireless channel summaries were combined within each session before descriptive aggregation across sessions. Quiet-window EEG RMS reflected both physiological activity and recording noise and was interpreted separately from input-short noise.

### PTZ experiments and seizure annotation

PTZ was administered at 40 mg kg^−1^ after baseline recording. Baseline analyses used accepted pre-administration data.

Three experienced researchers independently reviewed multichannel recordings for electrographic seizures, visually defined as distinct repetitive polyspike discharges exceeding twice baseline amplitude and 5 Hz for longer than 5 s. An interval was accepted when at least two reviewers judged that it met these criteria. Seizure onset and offset marked the beginning and termination of the qualifying sustained discharge. Recordings with insufficient analyzable duration or severe multichannel signal loss were excluded.

For Fig. 3, repeated baseline segments and seizures were summarized within each animal. The PSD embedding included 13 accepted segments and 1,035 windows from four animals. State comparisons used baseline and seizure categories without separate preictal or postictal labels.

### Seizure detector development and evaluation

The detector represented a 30-s history of six cortical channels as 27 overlapping 4-s windows advanced in 1-s steps. Preprocessing included timestamp regularization to 1,000 Hz, demeaning, 0.1-Hz high-pass filtering, and 50-Hz notch filtering. During training and retrospective evaluation, each 4-s window was extracted from its own preceding filter history of up to 30 s, with resampling from 1,000 to 500 Hz and then to 256 Hz. During deployment, the latest 30-s buffer was filtered once and resampled directly to 256 Hz before constructing the 27 windows, reducing repeated preprocessing. Both implementations applied forward–reverse filtering to available historical data only. Within each context, channel-wise normalization used the median and the 95th percentile of absolute deviations across windows and time points.

A shared ShallowConvNet encoder mapped each 4-s window to a 16-dimensional token^40^. A single-layer GRU with 16 hidden units integrated the sequence of 27 tokens, and a shared classifier generated a prediction at each time step^41^. The model contained 53,434 trainable parameters. Seven architectures were evaluated across 10 random seeds: ShallowConvNet–GRU, EEGNet–GRU, ShallowConvNet, DeepConvNet, STFT-small-CNN, PSD-CNN and PSD-MLP^40,42,43^. STFT denotes short-time Fourier transform, CNN convolutional neural network, and MLP multilayer perceptron.

Contexts were allocated chronologically to training, validation, and test subsets, comprising 189, 62 and 73 contexts, respectively. This subject-dependent training protocol included the same animals and recordings in multiple subsets, with filtering histories overlapping split boundaries. Evaluation therefore assessed detection in later portions of the same recordings.

The selected ShallowConvNet–GRU was trained for 130 epochs without early stopping, using AdamW with a learning rate of 0.001, weight decay of 10^−5^, and a batch size of 64. Training used class-weighted cross-entropy, with the positive-class weight multiplied by 0.7, and gradient clipping at a maximum norm of 5. Checkpoints were selected by validation window-level F1. The decision threshold was selected on the validation subset and then fixed for testing.

Performance was evaluated separately at the window and event levels^44^. Window-level precision, recall, and F1 pooled predictions from all 27 internal 4-s windows in each 30-s test context, yielding 1,971 overlapping predictions across 73 contexts. These predictions were scoring units, not independent recording or seizure replicates. Streaming inference retained the final sequence output as the current seizure probability, which the GUI updated once per second.

For any-overlap (OVLP) scoring, recall was the proportion of reference events overlapping at least one prediction for a nonzero duration, and precision was the proportion of predictions overlapping at least one reference event^45^. Neither full-event coverage nor one-to-one matching was required. Time-aligned event scoring (TAES) used a continuous-time adaptation of the Neural Engineering Data Consortium (NEDC) Eval EEG procedure^45^. It awarded partial credit for reference-event coverage and penalized predictions extending beyond reference boundaries; unmatched references and predictions counted as misses and false alarms, respectively. Exact calculations and rules are provided in Supplementary Methods S2.

For each scoring definition, F1 was calculated as:

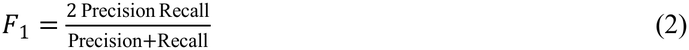

For pooled window classification, precision = TP/(TP + FP) and recall = TP/(TP + FN), with counts pooled across the 1,971 internal 4-s test-window predictions. Event-level precision and recall followed their respective scoring rules; Equation (2) was applied separately to each evaluation. Pooled window metrics summarized all internal predictions; deployment retained the latest sequence output at each 1-s update.

Software runtime was measured in three HDF5 pseudo-online replays on an NVIDIA RTX 4090, each comprising 520 updates. CUDA synchronization was performed before and after model execution. Model forward-pass timing excluded preprocessing, input preparation, device transfer and post-processing. Software processing time spanned submission of a complete 1-s data chunk to return of the prediction result, including buffer updates, signal-quality checks, preprocessing, model execution, and output processing. It excluded disk access, Bluetooth transmission, acquisition waiting, queueing and GUI rendering.

For both latency measures, timings were averaged within each replay, and the reported mean ± SD summarized the three replay means. A separate controlled comparison of model architectures used a batch size of one with inputs already resident on the GPU. These measurements excluded initial-context accumulation and did not quantify the delay from seizure onset to detection.

### Offline cortical signal analyses

Representative Morlet maps used the pointwise median across six robustly normalized channels. Wavelet power over 1–100 Hz was expressed in dB relative to each frequency’s temporal median within the displayed segment.

For band-power analysis, baseline and seizure intervals were divided into 4-s windows advanced by 2 s. Median-centered signals underwent Welch PSD estimation using 2-s subsegments with 50% overlap^39^. PSD was integrated over delta (1–4 Hz), theta (4–12 Hz), beta (13–30 Hz), low gamma (30–55 Hz) and high gamma (55–100 Hz).

Channel powers were converted to dB and averaged across channels. Window medians were then calculated within events or segments, followed by medians across events or segments within animal and condition.

The displayed cross-channel band-power summary was:

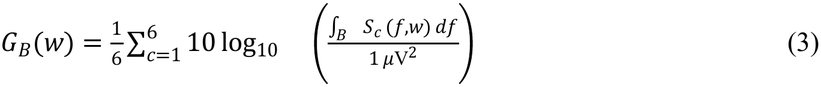

Here B denotes the frequency band, w the analysis window and *S_c_* (*f,w*) the PSD of channel c in μV^2^/Hz. The reference was 1 μV². Averaging channel powers after conversion to dB corresponds to the logarithm of their geometric mean in linear units, rather than converting the arithmetic mean power to dB.

The PSD embedding used 10-s windows advanced by 5 s. Each channel’s spectrum was pooled into 100 bins over 0.5–100 Hz and normalized to unit sum. The six spectra formed 600 features, which were standardized and reduced by principal component analysis (PCA), retaining at least 95% of the variance. Euclidean t-SNE used perplexity 30^46^; k-means used two clusters and 50 initializations. Cluster-to-state matching was performed post hoc for exploratory visualization, without held-out classification.

PLV and ordinary amplitude-weighted wPLI were calculated for all 15 cortical pairs in 4-s windows advanced by 2 s, using band-pass filtering and Hilbert analytic signals. Bands were theta (4–12 Hz), beta (13–30 Hz), low gamma (30–55 Hz) and high gamma (65–100 Hz). Window estimates were aggregated through events or segments to animal and condition. Group maps displayed the median animal-wise seizure-minus-baseline difference. For each metric, the displayed band maximized the sum of absolute group changes, and the eight edges with the largest absolute changes were retained. Display selection was based on effect magnitude.

Within each window, PLV was calculated from the analytic-signal phases^23^:

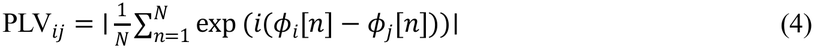

Here *ɸ_i_*[*n*] and *ɸ_j_*[*n*] are the Hilbert phases of channels i and j, and N is the sample count. For the corresponding analytic signals *z_i_* and *z_j_*, let I_ij_[n] = Im(z_i_[n] z_j_[n]*), where * denotes complex conjugation. Ordinary amplitude-weighted wPLI was^24^:

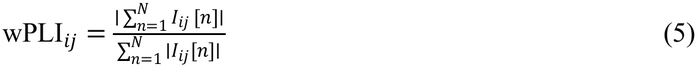

Equation (5) defines ordinary wPLI rather than debiased squared wPLI. Estimates were treated as missing when the mean absolute imaginary cross-product was ≤10^−12^.

Sharp transients were detected from absolute 5–70 Hz filtered signals. For each animal and channel, the baseline threshold was the median absolute amplitude plus 5 × 1.4826 × median absolute deviation (MAD). Peaks were separated by at least 50 ms and required an above-threshold duration of 20–200 ms. Rates were expressed as counts per minute and summarized by the median across segments within each animal and condition.

### Behavioral tracking and neural coupling

Overhead RGB videos were analyzed with SLEAP to track the nose, head, upper trunk, lower trunk and hip^47^. Body position was derived from valid trunk landmarks, converted from pixels to centimeters using arena-corner calibration, and smoothed before calculating displacement-based speed.

In the 40 × 40 cm open field, speeds below 0.5 cm s^−1^ were classified as stillness and speeds at or above this threshold as movement. Recordings were divided into 1-s windows advanced by 0.5 s, retaining windows with at least 90% state purity. Within each session and state, auto- and cross-spectra were pooled across accepted windows and discrete prolate spheroidal sequence (DPSS) tapers^48^ (NW = 2; three tapers).

Magnitude-squared coherence was then calculated as:

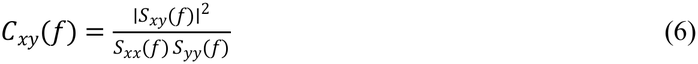

Here *S_xy_* is the cross-spectrum and *S_xx_* and *S_yy_* are the auto-spectra, averaged across accepted window–taper observations within each session and state before taking the ratio^49^. Line-noise bins were excluded from band estimates. Statistical tests used the inverse hyperbolic tangent of the square root of each session’s band-averaged coherence:

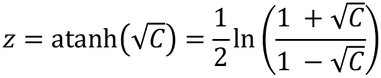

Here C denotes the session-level band-averaged coherence, clipped to [0, 0.999999] before transformation to avoid an infinite value at C = 1.

Transformed session estimates were averaged within each mouse (n = 4) and state before paired comparisons. Network maps instead displayed moving-minus-still differences calculated from the corresponding untransformed coherence estimates.

Social contact was defined as a minimum distance of ≤2 cm between the two animals’ tracked body points. Recordings were divided into 0.5-s windows advanced by 0.25 s, retaining windows with at least 90% state purity. Within each session, windows with the same label were grouped into an event when consecutive window starts were separated by no more than 0.5 s. Window-level features were averaged within each event. For each mouse and condition, event means were pooled across available sessions and averaged with equal weight, yielding one contact and one non-contact value per mouse (n = 5).

### Visual stimulation and response analysis

Right visual cortex recordings during visual stimulation were analyzed for periodic components near the 5.25-Hz reference frequency and its harmonics. Spectral analyses used the full preprocessed recording; waveform visualization used a selected continuous excerpt from the same recording. Welch PSD was estimated using 8-s Hann windows with 50% overlap^39^. Spectral parameterization with specparam/FOOOF separated periodic peaks from the aperiodic background^27^. Periodic excess was defined as:

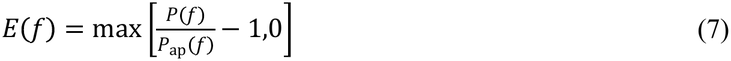

where P(f) is the measured PSD and *P*_ap_(*f*) is its fitted aperiodic component. Harmonic SNR was calculated as 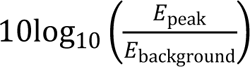, where *E*_peak_ was the maximum periodic excess within ±0.05 Hz of each reference harmonic, and *E*_background_ was its median within ±0.25 Hz after excluding the peak interval.

For waveform visualization, 80 non-overlapping windows were extracted from the selected excerpt. Each window spanned five cycles of the 5.25-Hz reference period, approximately 0.952 s. Windows were defined by the reference period, without alignment to measured stimulus triggers or designation as individual stimulation trials.

### ECG recording and analysis

ECG was recorded using the same acquisition electronics and wireless workflow, with electrodes placed subcutaneously near the right forelimb and left hindlimb. The differential signal was calculated as channel 8 minus channel 1 and filtered using a third-order Butterworth band-pass filter at 3–40 Hz, applied forward and backward. Beat detection and interval analysis used a selected stable segment spanning 80–560 s after recording onset.

Prominent negative-going QRS deflections were detected as peaks in the inverted signal, with a minimum separation of 75 ms and a minimum prominence of three times a robust amplitude scale, calculated as the median absolute deviation from the signal median divided by 0.6745. Beat intervals were calculated from the sample-index differences between successive retained detections, divided by the estimated sampling rate. Only intervals of 75–200 ms were included in the interval distribution and instantaneous heart-rate analysis. Heart rate was calculated as 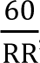, with RR expressed in seconds.

For waveform visualization, 30 eligible cardiac cycles from a representative segment were aligned to the detected QRS fiducial points. Each cycle extended from 45 ms before to 70 ms after the fiducial point, and aligned waveforms were summarized by their mean and sample standard deviation.

### Statistical analyses

Analyses were performed in Python using NumPy, pandas and SciPy. Descriptive units were channels and windows for electronic-noise summaries, recording sessions for commercial-system comparisons, and animals for PTZ and behavioral summaries. Model-performance variability was summarized across training seeds, and timing variability across replay runs. Repeated channels, windows, events, beats and sessions were not counted as independent animal replicates. PTZ and commercial-system comparisons were descriptive, and model architectures were compared without formal hypothesis testing.

Movement and social-contact effects were assessed using two-sided paired t-tests on animal-level estimates (n = 4 and n = 5 mice, respectively). Movement analyses used the transformed coherence values defined above, whereas social-contact analyses used the untransformed bandwise spectral coherence index. For each analysis and frequency band, Benjamini–Hochberg correction^50^ was applied to the 15 unique cortical-pair comparisons. Thus, each set of 15 tests formed a separate correction family; P values were not pooled across frequency bands or between the movement and social-contact analyses. Results were reported as nominally significant at unadjusted P < 0.05 or FDR-significant at adjusted P < 0.05.

## Data availability

The data supporting the findings of this study are being curated and will be made publicly available in a suitable repository. Repository details and persistent identifiers will be added to an updated version of this preprint.

## Code availability

The acquisition software, host application, analysis scripts, and seizure-detection code are being organized for public release. Repository details will be added to an updated version of this preprint.

## Acknowledgements

The authors thank the Tsinghua University Student Research Training Program, the Xianfeng Program of Weixian College, and the Future Instrument Student Team (FIST) of the Department of Precision Instrument, Tsinghua University, for their support.

We thank Rosa Chan, Xinge Yu, Chengqiang Tang, Zuowei Sun, Weixuan Huang and Xinying Cai from City University of Hong Kong for reviewing the manuscript and providing valuable comments and suggestions. We further thank Yiqun Wang, former president of FIST, and Haoren Xue, its current president, for their support. We also thank Mingrui Wang, Weihao Zhao, Yu Wu, Han Jiang, Wenrun Song, Yangxuan Zhou, Shuran Liu, Lin Gao, Chui Kong, and Jingyi Qiu for their assistance with the experiments, and Jianping Yu and Ru You for their support in laboratory management.

## Author contributions

W.Z. designed the acquisition circuitry, electrodes, and head-mounted mechanical components; developed the firmware, Bluetooth communication, and host application; designed the experiments; performed animal surgeries and behavioral experiments; characterized electronic noise and packet loss; compared recordings with the PowerLab system; conducted cortical coherence and phase-coupling analyses; prepared Figs. 1, 2, and 6; and wrote the original manuscript.

Y.D. developed and evaluated the seizure-classification model; analyzed seizure recordings; performed runtime benchmarking; contributed to SLEAP-based behavioral tracking and movement and social-contact annotation; prepared Figs. 3 and 4; and contributed to manuscript writing.

J.L. performed animal surgeries, behavioral experiments, and visual-stimulation experiments, and analyzed ECG recordings. Y.D. and J.L. curated the experimental data. Y.D., J.L., and H.W. independently annotated electrographic seizures.

G.L. analyzed the visual-stimulation recordings and prepared Fig. 5. C.Y. performed behavioral experiments and contributed to SLEAP-based behavioral tracking and movement and social-contact annotation. M.C. performed ECG electrode implantation and acquired ECG recordings.

X.X. and H.D. contributed to data analysis and reviewed and edited the manuscript. H.W. reviewed and edited the manuscript.

S.S., Y.Z., and Z.M. provided scientific guidance and reviewed and edited the manuscript.

## Supplementary information

Supplementary Information accompanies this paper and comprises Supplementary Figs. S1–S7, Supplementary Methods S1–S5, Supplementary Table S1 (analysis-specific preprocessing) and Supplementary Table S2 (comparison with commercial electrophysiology systems).

